# The Maryland Analysis of Developmental EEG (MADE) Pipeline

**DOI:** 10.1101/2020.01.29.925271

**Authors:** Ranjan Debnath, George A. Buzzell, Santiago Morales, Maureen E. Bowers, Stephanie C. Leach, Nathan A. Fox

**Author notes:** **Correspondence:** Ranjan Debnath, The Child Development Lab, Department of Human Development and Quantitative Methodology, University of Maryland, College Park, 20742 MD.

## Abstract

Compared to adult EEG, EEG signals recorded from pediatric populations have shorter recording periods and contain more artifact contamination. Therefore, pediatric EEG data necessitate specific preprocessing approaches in order to remove environmental noise and physiological artifacts without losing large amounts of data. However, there is presently a scarcity of standard automated preprocessing pipelines suitable for pediatric EEG.

In an effort to achieve greater standardization of EEG preprocessing, and in particular for the analysis of pediatric data, we developed the Maryland Analysis of Developmental EEG (MADE) pipeline as an automated preprocessing pipeline compatible with EEG data recorded with different hardware systems, different populations, levels of artifact contamination, and length of recordings. MADE uses EEGLAB and functions from some EEGLAB plugins, and includes additional customizable features particularly useful for EEG data collected from pediatric populations.

MADE processes event-related and resting state EEG from raw data files through a series of preprocessing steps and outputs processed clean data ready to be analyzed in time, frequency, or time-frequency domain. MADE provides a report file at the end of the preprocessing that describes a variety of features of the processed data to facilitate the assessment of the quality of processed data. In this paper we discuss some practical issues, which are specifically relevant to pediatric EEG preprocessing. We also provide custom-written scripts to address these practical issues.

MADE is freely available under the terms of the GNU General Public License at https://github.com/ChildDevLab/MADE-EEG-preprocessing-pipeline.

## Introduction

Electroencephalography (EEG) provides a measure of human neural activity with a high degree of temporal precision and the ability to characterize neuronal oscillations (Cohen, 2014; Luck, 2014; Nunez & Srinivasan, 2006). There has been a recent surge of interest in the use of EEG as a reflection of brain activity, particularly for research in pediatric populations. For example, recent work in human infants has shown how specific measures of EEG might be useful for characterizing the effects of prenatal experience, early life adversity, as well as identifying infants at risk for different developmental disorders (Marshall, Fox, & Group, 2004; Orekhova et al., 2014). Unlike EEG data recorded from adults, however, EEG recorded from pediatric populations is particularly susceptible to artifact contamination and only short recording sessions can be tolerated. Thus, there is concern about the amount of artifact-free EEG that can be acquired from pediatric populations. Moreover, the traditional frequency bands of EEG (e.g., delta, theta, alpha, beta) are not defined in the same way for younger participants, compared to adults (Marshall, Bar-Haim, & Fox, 2002). For example, the peak frequency for oscillations within the alpha band changes over the first years of life, which may further necessitate specific preprocessing decisions when analyzing pediatric data.

Before analyzing EEG to calculate neural measures of interest, it is necessary to perform a set of preprocessing steps (Luck, 2014), which serve to remove environmental noise and physiological artifacts. While there is general consensus as to what needs to occur during EEG preprocessing, the exact preprocessing steps vary amongst research labs, and as noted above, may differ for pediatric data. Moreover, a number of these preprocessing steps require subjective inputs and decisions by the user, which can result in further variability within and across labs. In an effort to achieve greater standardization of EEG preprocessing, particularly for the analysis of pediatric data, the Child Development Lab at the University of Maryland developed a preprocessing pipeline to exclude unwanted artifacts from data and improve the signal-to-noise ratio while minimizing data loss. The Maryland Analysis of Developmental EEG (MADE) pipeline achieves complete automation of EEG preprocessing, allowing objectivity and reproducibility, which is particularly well-suited for large-scale, multi-site projects.

While there are other publicly available EEG preprocessing pipelines (Bigdely-Shamlo, Mullen, Kothe, Su, & Robbins, 2015; Mognon, Jovicich, Bruzzone, & Buiatti, 2011; Nolan, Whelan, & Reilly, 2010), the majority of these pipelines are optimized for adult EEG data. Although, recently HAPPE preprocessing pipeline has been optimized for pediatric populations, it is not suitable for preprocessing data intended for event-related potential (ERP) analyses (Gabard-Durnam, Mendez Leal, Wilkinson, & Levin, 2018). The MADE pipeline was created to provide a variety of improvements over existing pipelines. First, MADE can process both resting state and event-related data from multiple different recording systems. Second, the MADE pipeline is transparent, as it is a set of MATLAB scripts that can be examined by the user, and is easily customizable by setting a few parameters at the beginning of the script. Third, the MADE pipeline utilizes advanced and automated artifact detection and correction procedures including a newly developed set of routines to modify current ICA approaches (Leach et al (under review). Finally, the MADE pipeline provides a variety of supplemental scripts to assist with re-labeling event-related data and excluding interference trials from infant data.

In line with the principles of open science, we have made our scripts publicly available (https://github.com/ChildDevLab/MADE-EEG-preprocessing-pipeline). This manuscript serves as a companion paper to the set of publicly available scripts and achieves two purposes: 1) it provides a detailed treatment of the theory behind each of the preprocessing steps implemented, so that novices and experts alike can understand the rationale behind each preprocessing decision, and 2) it shows how the pipeline performs compared to other preprocessing approaches. Additionally, the online supplement of this manuscript provides a step-by-step tutorial of how to use and customize the available scripts so that users, even with little-to-no knowledge of EEG data analysis, can utilize the MADE pipeline. Our goal is to provide a simple, user-friendly, and effective EEG data-preprocessing pipeline to facilitate research with pediatric EEG data.

### Pipeline overview

Our preprocessing pipeline is implemented as a set of MATLAB (The MathWorks, Natick, MA) scripts, which allow for complete automation of EEG preprocessing. The scripts that make up the pipeline draw heavily on the EEGLAB toolbox (Delorme & Makeig, 2004) functions and rely on the EEGLAB data structure as an organizing principle. We further leverage specific functions from some EEGLAB plugins for the identification of bad electrodes and artifactual independent components, respectively. The pipeline also includes additional customized features particularly useful for EEG data collected from pediatric populations, such as trial-level channel interpolation (e.g., Buzzell et al., 2019). Thus, the MADE pipeline reflects a novel combination of current state-of-the-art in EEG processing techniques, well-suited for pediatric data.

The steps involved in our preprocessing pipeline (Figure 1) include: high and low-pass filtering, automated identification/removal of bad EEG channels, independent components analysis (ICA) to identify and remove artifacts, creating epochs, artifact rejection on epoched data using voltage thresholding, channel interpolation, and re-referencing of epoched data. In the section below we describe the theory behind our specific approach to each of these preprocessing steps.

**Figure 1.**
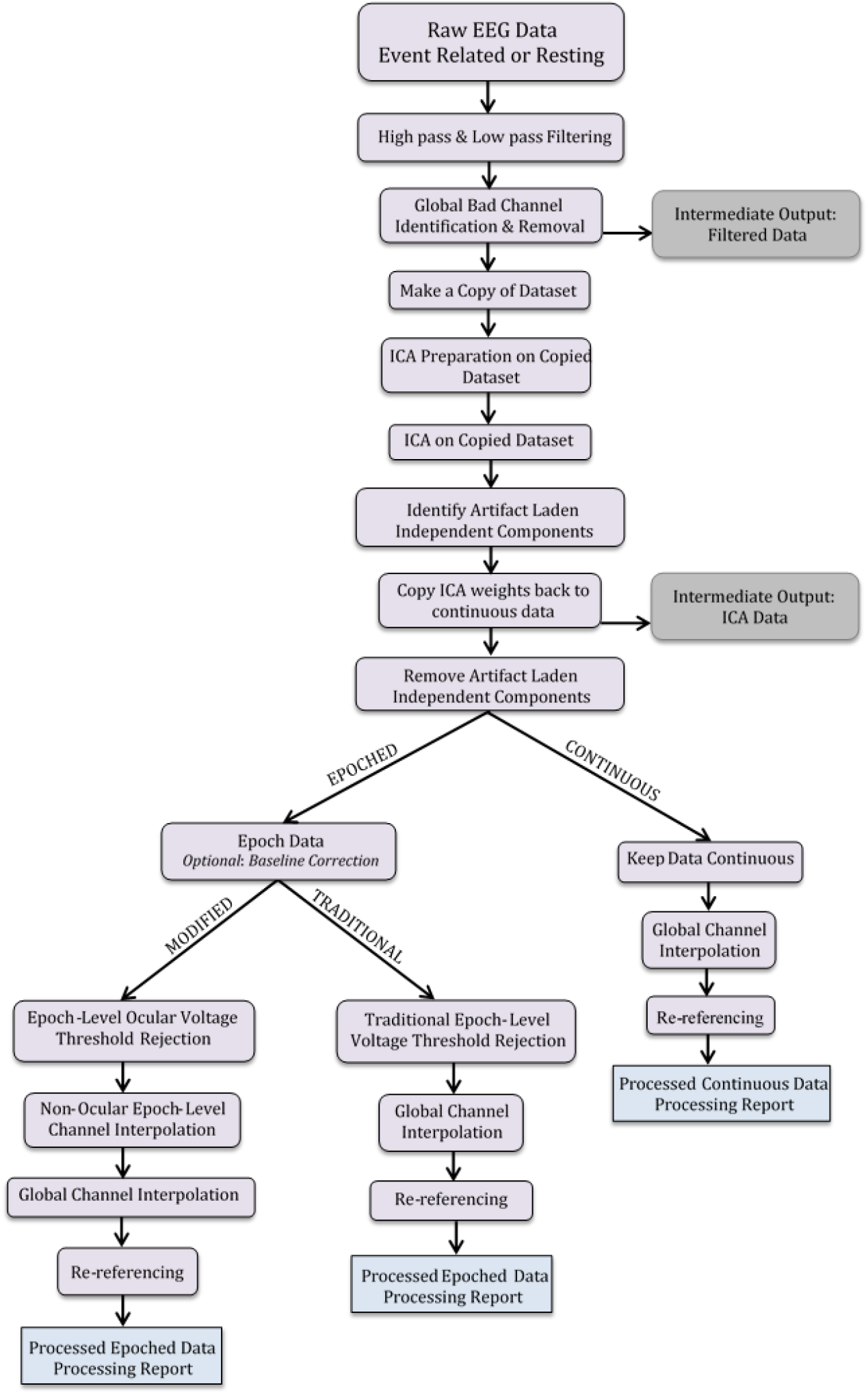
Schematic representation of MADE pipeline’s preprocessing steps. Independent component analysis is abbreviated to ICA. The intermediate results are indicated by the suffix added to the file name in that specific processing step in the gray boxes.

#### Filtering the data

Filtering EEG data can remove low-frequency drifts, skin potentials, high-frequency noise, EMG artifacts and electrical line noise that commonly manifest at 50Hz or 60Hz. We high-pass filter our data at .3 Hz and low-pass filter at/below the frequency of electrical line noise (50Hz or 60Hz), which is recommended for typical experiments studying cognitive, affective, and perceptual processes (Luck, 2014). We have tried several procedures to remove electrical line noise and found them unsatisfactory; therefore, as part of our standard preprocessing stream, we prefer filtering out frequencies at/above electrical line noise (50 Hz or 60 Hz). To avoid a latency shift that can be caused by some filters, we use a noncausal Finite Impulse Response (FIR) filter using the with FIRfilt plugin of EEGLAB (developed by A. Widmann: www.unileipzig.de/~biocog/content/widmann/eeglab-plugins/). Furthermore, we filter our continuous data (before creating epochs) to avoid edge artifacts in each data epoch. To minimize additional artifact that is created by a “steep” filter rolloff (see Luck, 2014), our low-pass filter is designed to have a 10 Hz transition band; we further apply a high-pass filter with a passband of .3 Hz and a stopband of .1 Hz. The reason for employing a passband of .3 Hz for the high pass filter, as opposed to .1 Hz, is because empirical research has shown that data recorded via high impedance systems (e.g. the EGI system) are susceptible to ultralow frequency artifacts (e.g. skin potentials) that need to be filtered out (Kappenman & Luck, 2010). We prefer applying the high and low-pass filters before identifying bad channels since filters can minimize noise and therefore improve detection of bad channels.

#### Removal of bad channels

Electrodes can “go bad” during the recording of EEG data for a number of reasons, including displacement due to head or body movements, changes in impedance, or faulty wiring. Pediatric data are often more susceptible to the issues of displacement or impedance change given that children are less likely to remain still and are often less tolerant of electrode adjustments (e.g., moving hair or re-seating electrodes) after capping. Therefore, it is common to identify bad channels during preprocessing of EEG data, particularly for pediatric data.

In order to detect bad channels, we use the ‘channel_properties.m’ function from the FASTER EEGLAB plugin (Nolan et al., 2010). The ‘channel_properties.m’ function identifies bad channels by first measuring three values that are standardized across all electrodes: the Hurst exponent, correlation with other channels, and channel variance. The Hurst exponent refers to a measure of the long-range dependence of time series data, with human EEG known to have a Hurst exponent of ~ 0.7; prior work has shown that deviations from a Hurst exponent of 0.7 can be used to detect the presence of a non-biological signal (Nolan et al., 2010). Therefore, if the Hurst exponent for a given channel is abnormal, then it is likely that the channel contains data that is primarily non-biological in nature (e.g. environmental noise). The correlation measure assesses how similar a given channel’s data is to other nearby channels, with the assumption that nearby channels should not be identical but should still be similar as a result of volume conduction (Nolan et al., 2010). Thus, if a given channel’s correlation value is abnormal, the channel likely did not record primarily brain activity. Finally, channel variance measures how variable the data for a given channel is over time; if the channel variance is abnormal, then it is assumed that the channel likely did not record primarily brain activity. The ‘channel_properties.m’ function measures the Hurst exponent, correlation, and channel variance values for all channels and then standardizes them by transforming into Z-scores. We consider any channel that has an absolute Z-score greater than 3, for any of the three measures, to be considered a bad channel. We delete bad channels globally (across the entire recording period) and these channels are interpolated at a later step (described in detail below). We recommend not deleting more than 10% of the channels for a particular participant.

#### Optional removal of outer electrodes for infants’ data (a priori)

One of the benefits of the EGI geodesic net, which is commonly employed in pediatric EEG research, is that it provides extensive coverage of the head (Tucker, 1993). However, we have found that the outermost ring of electrodes that are located near the base of the skull tend to have poor connections and are often noisy when recording from infants. For this reason, our standard procedure for preprocessing infant data is to remove this outer ring of electrodes, a priori, before running the ‘channel_properties.m’ function. The reason for removing these electrodes a priori, instead of relying on the ‘channel_properties.m’ function to identify and remove them, is because this function operates via calculating standardized values. Therefore, if a large number of bad channels are present in the data, then this will reduce the Z-score values calculated by the ‘channel_properties.m’ function, potentially making it more likely that not all bad channels are detected. This problem can be mitigated if a set of known bad channels (i.e. the outer ring of electrodes) is first removed.

#### Independent components analysis (ICA)

Even after filtering and the removal of bad channels (detected using either the FASTER tools or a priori removal described above), EEG recordings still contain a number of non-neural artifacts, including electrical deviations caused by: blinks, saccades, or EMG (Luck, 2014). One option for dealing with such artifacts is to simply identify the time segments in the data during which such artifacts are present and remove these segments completely from further analysis. This approach is valid and provides strong protection against misinterpreting physiological artifacts as neural data. However, in order to employ such an approach, it is necessary to either record data from a participant that is able to minimize blinks, saccades and muscle movements, or have a large amount of data so that even after throwing out such segments there is still an adequate amount of data left to analyze. Unfortunately, both of these options are often not feasible when analyzing EEG data from pediatric populations, especially infants. Therefore, an alternative approach to dealing with artifacts is to use ICA (Jung et al., 200; Delorme & Makeig, 2004) to identify such artifacts and then subtract the artifact-related activity from the rest of the EEG signal. ICA has the benefit of retaining the segments of data during which the artifacts occurred and is the approach that we employ in our pipeline.

ICA has been shown to perform better on data that retains a degree of stationarity (Winkler, Debener, Müller, & Tangermann, 2015). For example, ICA performs better when the data is first filtered with a 1 Hz highpass filter and periods of the recording that contain large amounts of EMG or periods where electrodes exhibit unusually high/low amplitudes (e.g. +/− 1000 μV) are additionally removed (Viola, Debener, Thorne, & Schneider, 2010; Winkler et al., 2015). However, an issue with applying a 1 Hz filter and removing segments of data with EMG before running ICA is that researchers may be interested in low-frequency information or time periods during which excessive EMG occurs. Therefore, a hybrid approach involves making a “copy” of the data, applying a 1 Hz high-pass filter and removing segments containing excessive EMG or high/low amplitude data from the copy, running ICA on the copy, then copying the ICA weights (which contain the information needed to identify artifacts) back to the original dataset that has not been filtered with the 1 Hz high-pass filter (Viola et al., 2010). However, when removing segments that contain excessive EMG or unusually high/low amplitudes in the copied dataset, it is possible that a few bad channels (missed by the FASTER tools described above) lead to the removal of too many data segments prior to running ICA on the copy. Therefore, prior to removing any segments of data from the copy, we first remove any channels that contain excessive EMG or unusually high/low amplitudes for greater than 20% of the recording; these same channels are removed from the original dataset as well. The end result of the copy/ICA process is an improved ICA decomposition without having to sacrifice low frequency information or time periods that contain excessive EMG or unusually high amplitudes.

Once the ICA decomposition has been completed, it is necessary to select independent components (ICs) that correspond to artifacts (blinks, saccades, EMG) and then subtract these components from the data. Several algorithms have been developed in order to automatically select ICA components, with one of the most popular being the ADJUST EEGLAB plugin (Mognon et al., 2011). This toolbox performs similarly to human observers when applied to adult data (Mognon et al., 2011). However, we found that ADJUST does not perform as well on pediatric data and misidentifies ICA components as artifact. Rather than manually reviewing the ICA components to correct the identification of ICA artifacts, we developed an alternative system. We modified the ADJUST scripts to improve ICA identification on pediatric data. Our “adjusted-ADJUST” scripts automatically select artifact laden independent components (ICs) and have been shown to perform better than the original ADJUST algorithm in adults, children, and infants (Leach et al., *under review*). After identifying ICs, we then subtract the ICA time series for these artifacts from the rest of the EEG signal. The final result is a continuous EEG data file with ICA-identified artifacts removed.

#### Epoching and removal of residual ocular artifact

To examine the task-related neural activity in EEG, it is common to cut the continuous EEG data into epochs (time segments) of data surrounding the experimental events before performing further analyses quantifying neural features within these epochs (Luck, 2014). Epochs are constructed by identifying event markers of interest and then cutting the continuous EEG data into epochs of appropriate length. For resting-state data, we recommend epoching; continuous resting-state data can be segmented into fixed length overlapping or non-overlapping epochs.

Once epochs have been created, it is possible to then loop through all epochs and identify any residual artifacts present within a given epoch. It is worth noting that the goal of removing ICA-identified artifacts is to “clean” the EEG signal without needing to completely reject time segments that contain ocular or other artifacts. However, we have found that ICA artifact identification and removal is rarely perfect, resulting in at least a small number of artifacts being missed and still present in the data following ICA-identified artifact removal. Therefore, we perform additional preprocessing steps to deal with such residual artifacts after epoching the data. First, we loop through each epoch and test whether the voltage recorded from a set of electrodes located near the eyes (ocular channels) exceeds a predetermined threshold (the exact threshold employed differs as a function of age, as discussed in the tutorial). If the voltage is exceeded at any one of the ocular channels for a given epoch, then we assume that residual ocular artifact is present within the epoch and reject the epoch (the epoch is removed from all further analyses). Next, we loop through all epochs a second time and identify whether any of the non-ocular channels (recorded from electrodes not located near the eyes) exceed the voltage threshold; for any epochs in which a non-ocular channel exceeds the voltage threshold, this channel is interpolated within that epoch using a spherical spline interpolation procedure (Perrin, Pernier, Bertrand, & Echallier, 1989). However, if greater than 10% of the non-ocular channels exceed the voltage threshold within a given epoch, then this epoch is instead rejected completely (removed from further analyses). To summarize our approach for dealing with residual artifacts: we use the electrodes located near the eyes to identify and reject any epochs that exceed a voltage threshold indicating the presence of residual ocular artifact, then additionally reject any epochs where more than 10% of the non-ocular channels exceed the voltage threshold; for all other epochs, individual channels are interpolated at the epoch level when they exceed the voltage threshold.

#### Channel interpolation

In the previous section, we noted that non-ocular channels exceeding a predefined voltage threshold during specific epochs are interpolated at the epoch level using a spherical spline interpolation procedure. However, even after these channels have been interpolated, there remain other channels missing from all epochs because they were rejected in one of the first preprocessing steps (those identified by the ‘channel_properties.m’ function). Following all other preprocessing steps, but before re-referencing the EEG data, we interpolate these missing channels using the spherical spline interpolation procedure as implemented in the EEGLAB toolbox. The reason for not interpolating these channels until all other preprocessing steps have been completed is because interpolated channels contribute no unique information to the ICA procedure (Delorme & Makeig, 2004). Additionally, the interpolated data will likely approach a closer estimate of the actual missing EEG data if other idiosyncratic artifacts are first removed from the channels being used to compute the interpolation.

#### Re-referencing

The last step we perform in our preprocessing pipeline is to re-reference the epoched data. Re-referencing means that the voltage time series for each electrode will no longer reflect voltage relative to the reference electrode(s) used during data collection. Instead, the voltage time series will now reflect voltage relative to offline, re-referenced electrode(s). Re-referencing is performed last, because in the case of computing an “average reference”, data from all channels are used in the computation of the average reference; therefore, all artifacts need to first be removed from all channels so that channel-specific artifacts are not propagated to all other channels during the computation of an average reference. Additionally, channel interpolation must be performed prior to computation of the average reference so that there is not a biased weighting of specific scalp locations in the estimate of the average reference.

### Validation Analysis of the MADE pipeline

In order to validate the MADE pipeline, we tested its performance on three datasets across childhood: an infant (12-month old) dataset, a childhood (3 – 6 years old) dataset, and a late adolescent (16-year old) dataset. We preprocessed 10 subjects in each dataset using three methods: A) the MADE pipeline; B) a traditional method involving only epoch level interpolation before artifact rejection, but no ICA-based artifact rejection (i.e., no FASTER and no ICA); and C) an even more conservative method without interpolation before artifact rejection. The outcome of interest was the percent of trials retained after each preprocessing method.

## Methods

Table 1 provides a description of the three EEG datasets used for testing the pipeline. In this section, we describe data acquisition procedures, the preprocessing steps and the results of the preprocessing of these three datasets.

**Table 1.** Acquisition parameters for 10 example data files from three age groups.

| EEG Dataset | Participant Age (years) | EEG type | EEG System | Sampling rate (Hz) | No. of channels | Reference |
| --- | --- | --- | --- | --- | --- | --- |
| Infant | 1 | Event Related | EGI | 500 | 128 | Cz |
| Childhood | 3-6 | Event Related | BioSemi | 512 | 64 | CMS |
| Late Adolescent | 16 | Resting | EGI | 500 | 64 | Cz |

### EEG data acquisition

#### Infant data

The infant dataset was part of a larger study examining the neural correlates of action observation and action execution in infants. Data were recorded with a 128-channel EGI system (NetAmps 300; Electrical Geodesics, Eugene, OR). The infants sat in their parent’s lap while viewing a variety of trials presented live by an experimenter behind a stage with a curtain. Infants saw 3 trial types in a block: observe grasp, observe point, and execute grasp. On each trial, the curtain was raised, and then the infant observed a colorful pinwheel spinning for 3 seconds as the baseline period. The curtain was then lowered before being raised again to show the live experimenter executing one of the three trials. Observe grasp trials involved the experimenter reaching for and grasping a toy in the middle of the stage. Observe point trials involved the experiment pointing to a toy. Execute grasp trials involved the stage being pushed toward the baby for the baby to reach and grasp the toy. There were 15 blocks of these three trial types in a randomized order and the infants completed as many trials as possible. The vertex (Cz) electrode was used as online reference. EEG data were sampled at 500 Hz and impedances were kept below 100 kΩ.

#### Child Data

The child data were part of a larger study examining the neural correlates of memory formation in children from a Northeast city from the United States (Riggins and Rollins, 2015). Data were recorded using a 64-channel BioSemi Active 2 EEG recording system for a study investigating age-related differences in memory (Riggins & Rollins, 2015). EEG data were recorded at 512 Hz and Common Mode Sense (CMS) electrode was used as reference. Children completed an ERP task in which they passively viewed items on a computer screen. There were three blocks and each block included 54 items that the children had been familiarized with the previous day and 27 new distractor items. Stimuli were presented on the screen for 500 ms, followed by a fixation cross that was jittered from 1,250 to 1,700 ms. Each block consisted of random presentation of the 54 previously seen (target items) and 27 new (distracter) items, for a total of 243 ERP trials.

#### Late Adolescent Data

The late adolescent dataset was part of a larger study examining the behavioral and neural effects of early neglect in children from Bucharest, Romania (Debnath et al., 2019). Data were recorded with a 64-channel EGI system (NetAmps 300) for a randomized, controlled trial of foster care placement in Bucharest, Romania (Debnath, Tang, Zeanah, Nelson, & Fox, 2019). The resting EEG was recorded for 6 min, alternating 1 min of eyes open and eyes closed. During the eyes open condition, the subjects were instructed to fixate on a small white cross in the center of a computer screen. The vertex (Cz) electrode was used as an online reference. EEG data were sampled at 500 Hz and impedances were kept below 50 kΩ.

### Preprocessing with MADE

EEG data from three studies were preprocessed with the fully automated version of the MADE pipeline and two more traditional methods (one with interpolation and one without). The following sections describe the MADE pipeline and the two traditional methods’ preprocessing steps that were applied to the three datasets and the post-processing reports generated by the MADE pipeline. In the infant dataset, EEG channels on the boundary of the electrode net were excluded from analyses since they were heavily susceptible to eye, face and head movements. This step removed 24 channels, leaving 104 channels included for further analysis. Infant and childhood data were down-sampled at 250 Hz. Continuous data collected with the EGI EEG system were high pass filtered at 0.3 Hz while the data collected with the BioSemi EEG system were high pass filtered at 0.1 Hz. The infant and child datasets were low pass filtered at 50 Hz and the adolescent dataset was low pass filtered at 40 Hz using FIR filters with a Hamming window with the FIRfilt plugin from EEGLAB. Artifact-laden channels were identified using FASTER (Nolan et al., 2010) and removed from analysis. To further remove ocular artifacts and generic noise, MADE performs independent component analysis (ICA). The ICA is performed on an identical copy of the dataset and the independent components (ICs) are transferred from the copied dataset to the original dataset. A copy of the dataset is made and further cleaned before performing ICA on this copied dataset. The copied dataset was high pass filtered at 1 Hz and segmented into 1s epochs. To achieve an improved ICA decomposition, noisy segments of data were rejected using a combined voltage threshold of ±1000 μV and spectral threshold (range −30 dB to +100 dB) within the 24–40 Hz frequency band to remove EMG-like activity. If this artifact rejection process identified more than 20% of the epochs for a given channel as containing artifact, that channel was removed from both the ICA copied dataset and the original dataset. After further cleaning, extended infomax (runica) ICA was performed on the copied dataset and the ICs were then transferred from the copied dataset to the original dataset. All further preprocessing steps were applied on this original dataset.

Individual components (ICs) containing artifacts were identified by an automatic process using the adjusted-ADJUST scripts (Leach et al., *under review*). The continuous EEG data were then segmented into fixed length epochs separately for each dataset. The infant data were epoched from −1s to 2s relative to the three event markers: execution grasp complete, observation grasp complete and observation point onset. The childhood data were epoched from −1s to 1s relative to the two event markers: target and distracter, and the late adolescent data were segmented into 2s epochs with 1s (50%) overlap. After segmenting data, the epochs in childhood data were baseline corrected using the time window from −1000 ms to −500 ms and the adolescent data were baseline corrected using the entire epoch (0–2000 ms), and the infant data were not baseline corrected. To further exclude artifacts from epoched data, a voltage threshold rejection (±150 μV) was applied in a set of frontal electrodes (infant data: E1, E8, E14, E21, E25, E32; childhood data: F7, F8, AF7, AF8, FP1, FP2, FPz; adolescent data: E1, E8, E25, E32). If an epoch in these frontal channels exceeded the voltage threshold of ±150 μV, that epoch was rejected. For all other channels, artifacted channels in each epoch were interpolated using the artifact free data from the surrounding channels within that epoch. If more than 10% of the channels within an epoch were interpolated, that epoch was rejected. After artifact rejection, missing channels were interpolated using spherical interpolation with the eeg_interp function from EEGLAB as implemented in the MADE pipeline. Epoched data were then re-referenced to an average of all channels. For each dataset, MADE produces a processing report for each EEG file in a single CSV file to evaluate preprocessing performance and data quality across subjects within a study. Table 2, 3, 4 present the processing report with all the metrics for the three example datasets. Figure 2 shows examples of EEG signals from the three datasets before and after MADE processing.

**Table 2.** MADE preprocessing report for the 10 files in infant dataset.

| EEG file | No. of recording channels | No. of preprocessed channels | No. of bad channels | EEG length (s) entered ICA | Total ICs | Number of bad ICs removed | Total epochs before artifact rejection | Total epochs after artifact rejection | Number of interpolated channels |
| --- | --- | --- | --- | --- | --- | --- | --- | --- | --- |
| p1201 | 128 | 104 | 1 | 1407 | 103 | 15 | 30 | 16 | 1 |
| p1202 | 128 | 104 | 0 | 1654 | 104 | 4 | 45 | 26 | 0 |
| p1203 | 128 | 104 | 3 | 1041 | 101 | 3 | 23 | 20 | 3 |
| p1204 | 128 | 104 | 1 | 1172 | 103 | 0 | 24 | 19 | 1 |
| p1205 | 128 | 104 | 4 | 1402 | 100 | 12 | 45 | 16 | 4 |
| p1207 | 128 | 104 | 1 | 1063 | 103 | 1 | 25 | 13 | 1 |
| p1208 | 128 | 104 | 2 | 956 | 102 | 19 | 24 | 20 | 2 |
| p1209 | 128 | 104 | 0 | 1071 | 104 | 11 | 24 | 17 | 0 |
| p1210 | 128 | 104 | 4 | 1500 | 100 | 0 | 41 | 33 | 4 |
| p1211 | 128 | 104 | 6 | 1177 | 98 | 1 | 20 | 15 | 6 |

**Table 3.** MADE preprocessing report for the 10 files in childhood dataset.

| EEG file | No. of recording channels | No. of preprocessed channels | No. of bad channels | EEG length (s) entered ICA | Total ICs | Number of bad ICs removed | Total epochs before artifact rejection | Total epochs after artifact rejection | Number of interpolated channels |
| --- | --- | --- | --- | --- | --- | --- | --- | --- | --- |
| 311 | 64 | 64 | 4 | 552 | 60 | 3 | 243 | 207 | 4 |
| 312 | 64 | 64 | 4 | 552 | 60 | 11 | 243 | 212 | 4 |
| 313 | 64 | 64 | 1 | 556 | 63 | 17 | 243 | 209 | 1 |
| 330 | 64 | 64 | 2 | 553 | 62 | 6 | 243 | 176 | 2 |
| 349 | 64 | 64 | 4 | 544 | 60 | 9 | 243 | 212 | 4 |
| 381 | 64 | 64 | 3 | 552 | 61 | 1 | 243 | 134 | 3 |
| 386 | 64 | 64 | 2 | 554 | 62 | 7 | 243 | 230 | 2 |
| 388 | 64 | 64 | 2 | 544 | 62 | 3 | 243 | 168 | 2 |
| 412 | 64 | 64 | 1 | 480 | 63 | 7 | 243 | 77 | 1 |
| 448 | 64 | 64 | 2 | 501 | 62 | 3 | 243 | 226 | 2 |

**Table 4.** MADE preprocessing report for the 10 files in late adolescent dataset.

| EEG file | No. of recording channels | No. of preprocessed channels | No. of bad channels | EEG length (s) entered ICA | Total ICs | Number of bad ICs removed | Total epochs before artifact rejection | Total epochs after artifact rejection | Number of interpolated channels |
| --- | --- | --- | --- | --- | --- | --- | --- | --- | --- |
| 005_eeg | 64 | 64 | 1 | 429 | 63 | 27 | 430 | 425 | 1 |
| 006_eeg | 64 | 64 | 3 | 415 | 61 | 11 | 414 | 414 | 3 |
| 008_eeg | 64 | 64 | 1 | 425 | 63 | 10 | 425 | 425 | 1 |
| 010_eeg | 64 | 64 | 1 | 424 | 63 | 15 | 423 | 423 | 1 |
| 011_eeg | 64 | 64 | 4 | 403 | 60 | 28 | 402 | 398 | 4 |
| 013_eeg | 64 | 64 | 2 | 429 | 62 | 14 | 430 | 430 | 2 |
| 014_eeg | 64 | 64 | 3 | 400 | 61 | 15 | 399 | 396 | 3 |
| 015_eeg | 64 | 64 | 5 | 450 | 59 | 12 | 452 | 386 | 5 |
| 016_eeg | 64 | 64 | 2 | 404 | 62 | 15 | 405 | 400 | 2 |
| 017_eeg | 64 | 64 | 4 | 396 | 60 | 6 | 400 | 391 | 4 |

**Figure 2.**
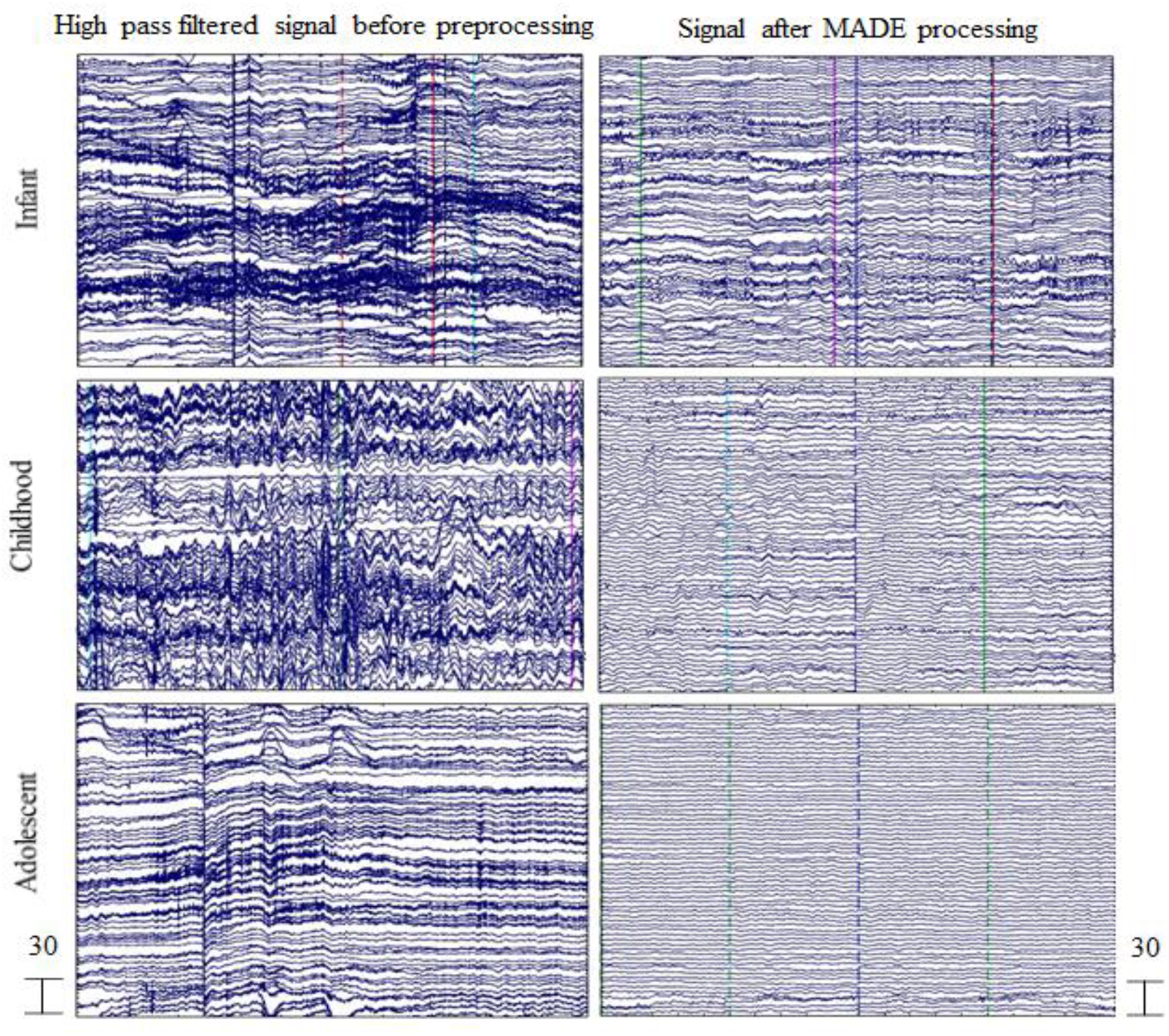
EEG signal before and after MADE processing. Three files from the three example datasets are shown with 5 s of data extracted from the recording. The EEG signal after high-pass filtering is shown in the left panel. The EEG signal after MADE processing as described in the preprocessing section of the example analysis is shown in the right panel. All scales are in microvolts.

### Traditional Preprocessing Methods

Similar to the MADE pipeline, the two traditional preprocessing methods applied a 0.3 Hz high pass filter to the continuous data collected with an EGI EEG system (0.1 Hz high pass for data collected with the BioSemi system) and a 50 Hz low pass filter to the infant and child datasets (40Hz low pass for the adolescent dataset). Infant and childhood data were down-sampled at 250 Hz. Next, the continuous EEG data were segmented into fixed length epochs separately for each dataset (using the same epochs for each dataset as the MADE pipeline). For the traditional method that included interpolation, a voltage threshold rejection (±150 μV) was applied in a set of frontal electrodes (the same sets the MADE pipeline used for each dataset). As in the MADE pipeline, the epochs in childhood data were baseline corrected using the time window from - 1000 ms to-500 ms, the adolescent data were baseline corrected using the entire epoch (0-2000 ms), and the infant data were not baseline corrected. If an epoch in these frontal channels exceeded the voltage threshold of ±150 μV, that epoch was rejected. For all other channels, artifacted channels in each epoch were interpolated using the artifact free data from the surrounding channels within that epoch. If more than 10% of the channels within an epoch were interpolated, that epoch was rejected. For the method that did not include interpolation, voltage threshold rejection, (±150 μV) was applied over all channels and any epochs where at least one channel exceeded this voltage threshold of ±150 μV were rejected. Finally, for both of the traditional preprocessing methods, the remaining epoched data were then re-referenced to an average of all channels.

## Results

The results of the comparisons of the three preprocessing methods are summarized in Table 5 and Figure 3. In order to compare the different methods, we performed Wilcoxon Signed Ranks tests on the proportion of trials retained for each preprocessing method, separately for the three datasets (Adolescent, Child, and Infant) as each dataset used a different task. To control for potential Type I errors due to multiple comparisons, we used the false discovery rate (FDR) correction (Benjamini and Hochberg, 1995). The FDR-corrected *p* value is represented by the *q* value.

**Table 5.** Descriptive Information on Trials Retained. Descriptive information on the proportion of trials retained by each preprocessing method (MADE Pipeline, Traditional with interpolation, and Traditional without interpolation) for each of the three datasets (Adolescents, Children, and Infants).

| Age |  | N | Mean | Std.<br>Deviation | Minimum | Maximum | Percentiles |  |  |
| --- | --- | --- | --- | --- | --- | --- | --- | --- | --- |
|  |  |  |  |  |  |  | 25th | 50th<br>(Median) | 75th |
| Adolescents | MADE Pipeline | 10 | 97.90% | 4.45% | 85.40% | 100.00% | 98.51% | 99.13% | 100.00% |
|  | Traditional with interpolation | 10 | 94.31% | 5.77% | 80.53% | 100.00% | 92.42% | 95.20% | 98.88% |
|  | Traditional without interpolation | 10 | 72.03% | 15.52% | 43.95% | 94.20% | 60.44% | 75.36% | 83.53% |
| Children | MADE Pipeline | 10 | 75.94% | 19.76% | 31.69% | 94.65% | 65.64% | 84.90% | 88.68% |
|  | Traditional with interpolation | 10 | 43.61% | 17.84% | 23.05% | 78.19% | 29.61% | 38.45% | 57.20% |
|  | Traditional without interpolation | 10 | 19.30% | 17.32% | 0.00% | 45.27% | 3.70% | 13.62% | 38.17% |
| Infants | MADE Pipeline | 10 | 67.44% | 16.86% | 35.56% | 86.96% | 53.00% | 72.92% | 81.20% |
|  | Traditional with interpolation | 10 | 31.84% | 11.53% | 13.33% | 48.00% | 22.46% | 31.88% | 41.46% |
|  | Traditional without interpolation | 10 | 21.70% | 11.26% | 6.67% | 41.67% | 11.88% | 20.42% | 31.50% |

**Figure 3.**
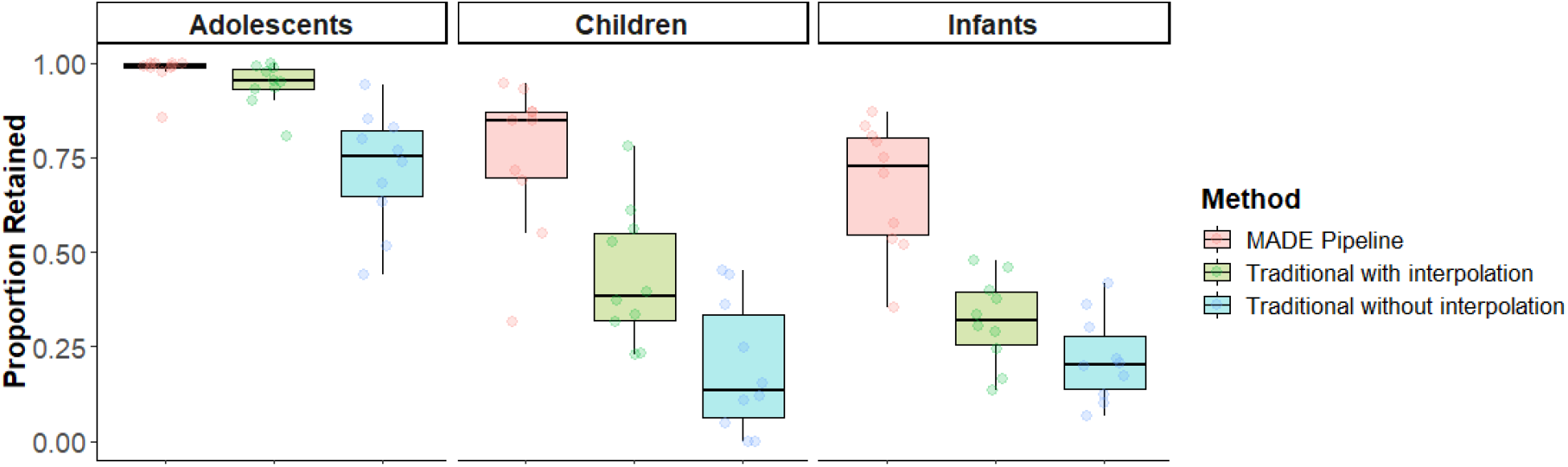
Box plots showing the proportion of trials retained by each preprocessing method (MADE Pipeline, Traditional with interpolation, and Traditional without interpolation) for each of the three example datasets (Adolescents, Children, and Infants) included in the analyses.

As seen in Figure 3, the MADE pipeline retained significantly more trials than the traditional preprocessing method with interpolation for adolescents, *Z*=2.52, *p*=0.012, *q*=0.012, children, *Z*=2.80, *p*=0.005, *q*=0.006, and infants, *Z*=2.80, *p*=0.005, *q*=0.006. The MADE pipeline also retained significantly more trials than the traditional method without interpolation for adolescents, *Z*=2.80, *p*=0.005, *q*=0.006, children, *Z*=2.80, *p*=0.005, *q*=0.006, and infants, *Z*=2.80, *p*=0.005, *q*=0.006. In sum, as expected, the MADE pipeline performed better than the two comparison methods.

## Discussion

Recent advancement in EEG data recording technology and the advent of high-density EEG nets have given impetus to using EEG in diverse population and large-scale studies. The use of EEG in large-scale studies with different subject groups yields large amounts of data. In many cases, the EEG data, particularly EEG data collected from pediatric populations, make the data processing procedure highly complex and challenging. The pediatric data often contain a high degree of artifact contamination, shorter recording lengths, and the quality of the EEG recordings is substantially reduced compared to recordings collected in adults. Traditionally, researchers relied on expert supervision for artifact identification and removal for pediatric EEG data. However, the manual data cleaning process decreases the replicability of methodology and the processed results. Moreover, with the increasing amount of available data and complexity of preprocessing procedures, the manual data processing becomes impractical and makes automatic preprocessing of EEG data essential. Multiple open source toolboxes and pipelines are publicly available for EEG data preprocessing (APP, Automagic, PREP) but these pipelines are optimized for data features that are often not particularly suitable for pediatric EEG data due to the constraints in pediatric EEG recordings. Although the HAPPE preprocessing pipeline has recently been streamlined to process pediatric data, it is unable to process ERP data. Therefore, there is a dearth of software with standard features used in adult EEG data processing (e.g. ICA, automatic identification of noisy channels) for processing of pediatric EEG data. To address these issues, we developed MADE, an open source MATLAB software that combines currently available standard preprocessing techniques and custom-made features particularly suitable for EEG data collected from pediatric populations.

We developed MADE as an automated EEG preprocessing pipeline for pediatric EEG data and investigated the effect of processing resting and task-based EEG data from diverse populations through MADE. This validation revealed that MADE performs significantly better than two other processing streams, which are the common data analysis approaches used in many labs. MADE retained significantly more trials compared to the traditional methods for both resting and task-based EEG data in all age groups. At end of the processing of a dataset, MADE produces a report file containing summary of quality measures for each datafile. The report file will allow the users to determine whether a particular subject needs to be excluded from analysis due to, for example, excessive contamination of artifacts or insufficient number of post-processed artifact free trials. MADE is also able to save data at different levels of the preprocessing pipeline, which would allow the users to examine whether a particular datafile might have some issue that needs a closer look at an earlier step of data processing. Moreover, MADE is able to process both resting and ERP data, which was a limitation in previous pipelines developed for pediatric data. Finally, MADE is very easy to customize for a particular dataset and users, we believe, will be able to seamlessly adapt MADE for their study.

There are some limitations to the MADE pipeline and the presented validation. We only tested MADE on 64- and 128-channel EEG datasets. However, we expect the pipeline to work equally well for recordings with more channels. We do not recommend the use of MADE for EEG data recorded with less than 32 channels. In such cases, MADE can still be used, but some parameter calculation and processing steps might not be optimal due to insufficient data channels. Furthermore, MADE relies heavily on ICA for artifact rejection, hence the correction of eye-movement related artifacts using EOG channels cannot be performed. However, the most challenging part of ICA based artifact rejection technique is to correct classification of ICs. In this pipeline, we used an automatic ICs classification method, the adjusted-ADJUST, to avoid subjective bias in ICs classification. The adjusted-ADJUST scripts were created by optimizing the ADJUST algorithm for infant data and are believed to be performing significantly better in pediatric populations than existing ICs classification methods. Finally, the validation of the pipeline and comparison of results with traditional methods are limited to the percent of post-processing trials. However, one of the main goals of this pipeline was to minimize loss of data, which is a major concern in EEG studies with pediatric population, while excluding unwanted artifacts from raw EEG signal.

In sum, this paper proposes and validates MADE, an EEG preprocessing pipeline streamlined for pediatric EEG data. We validated MADE on EEG data recorded with different systems and populations. Our results show that MADE performs significantly better than other traditional preprocessing methods. MADE is freely available under the terms of the GNU General Public License (version 3) (Free Software Foundation, 2007). MADE and associated scripts may be accessed at: https://github.com/ChildDevLab/MADE-EEG-preprocessing-pipeline. We hope that this automatic EEG data processing pipeline will contribute to pediatric EEG research and users will benefit from this pipeline and its accompanying MATLAB scripts.

## Supporting information

The online supplement of this manuscript provides a step-by-step tutorial of how to use and customize the available scripts.

## Appendices

## Appendix A. Excluding interference trials from analysis

In EEG experiments with pediatric populations, the subjects or caregivers often tend to make unintended motions in an experimental trial. The unintended motion can be of different natures depending on the experiment design such as gesture, gross motor movement, or leg movement. Moreover, there are trials in which the subjects do not perform the desired task event. The trials in which the subjects or caregivers appear to make unintended motions or the subjects do not perform the task are generally categorized as interference trials. In order to identify the interference trials, a common practice in pediatric EEG experiments is that the EEG tasks are video recorded and videos are coded for live events and synchronized with the continuous EEG recording. The coders view the videos off-line to identify the event of interest and also interference trials. The interference trials are then excluded from analysis.

There are different ways to code and exclude interference trials from analysis. Here we present a procedure, which is a standard method of our lab, to remove interference trials from EEG analysis. Two independent coders view each video file off-line frame-by-frame and separately identify each event of interest and all interference events. Coders identify the frame, in which an experimental event is completed and in which an interference event occurred. If the frame in which the interference occurred falls within an experimental trial, then that trial is coded as an interference trial to be excluded from analysis. After video coding, an excel file is created including the interference trials from all conditions and subjects. An example excel file of interference trials can be found on at https://github.com/ChildDevLab/MADE-EEG-preprocessing-pipeline.

To exclude the interference trials from analysis for each subject, we developed a method that includes a 3-step process: 1) reading the excel file containing interference trials of all subjects, 2) numerical labeling of all trials in each condition, 3) marking interference trials. The Matlab script exclude_interference_trials.m implements this method. The users can edit and adapt the script for their experiments and call the script to the pipeline for excluding interference trials from analysis.

## Appendix B. Segmenting eyes open and eyes closed resting state EEG data

Resting state EEG data are one of the most commonly recorded brain measures in pediatric populations. Because resting state EEG does not involve any specific task event, the data are segmented by inserting dummy markers. The EEGLAB function eeg_regepochs.m has been used in the MADE pipeline to segment a continuous dataset into consecutive epochs of a specified regular length by adding dummy markers and epoching the data around these markers. This way the whole length of continuous EEG data is converted into epochs. However, resting state EEG is generally also recorded with two conditions: eyes open and eyes closed. It is increasingly becoming a common practice to record resting state EEG for several minutes, alternating eyes open and eyes closed conditions. The eeg_regepochs.m function cannot be used to segment data into separate eyes open and eyes closed conditions. Therefore, we have developed a procedure to segment continuous resting EEG data into specific length epochs in the eyes open and eyes closed conditions. The Matlab script create_eyes_open_closed_resting_epoch.m provides a customized method of creating separate eyes open and eyes closed epochs from continuous EEG data recorded in eyes open and eyes closed conditions. The script takes users inputs for dummy markers and epoch length, then inserts dummy markers at specific intervals and creates epochs of the specified length that are time locked to those dummy markers. Furthermore, it can create either overlapping or non-overlapping epochs based on user’s inputs. The users can customize the create_eyes_open_closed_resting_epoch.m script to adapt it for their data.

## Appendix C. Marker editing for task-related EEG data

EEG files recorded while participants perform experimental tasks typically contain event markers that indicate when specific stimuli or motor responses were made. For example, a visual task requiring participants to indicate, via button press, what stimulus was presented to them will at least contain event markers indicating exactly when the stimuli were presented and button presses were made by the participant. Depending on what software was used to present the experimental task and what kind of EEG system was used to collect EEG data, the individual event markers may contain further information and/or additional markers that contain information about the task parameters. For example, if two kinds of stimuli are presented to participants within a single experimental task, then the stimulus identity may already be coded in the stimulus event marker, or, an additional event marker may contain information about the stimulus identity. Similarly, in order to determine whether a given stimulus event marker arises from an experimental trial in which the participant correctly responded, accuracy information from the response marker, or another marker, needs to be “copied over” to the associated stimulus event marker.

In order to properly label all event markers present in the EEG file, it is necessary for the MADE pipeline to call an additional script that loops through each event marker and properly labels them based on available information contained in other nearby markers. The script that will properly label the event markers in your EEG file needs to be customized for the purposes of the specific experimental task that is associated with your EEG file, as well as to be compatible with the software used to present the experimental task and EEG system used to collect the EEG data. As an example of what this script might look like, please refer to the script entitled, ‘edit_event_markers_example.m’. This script is a simplified version of a more complicated script that we commonly employ to label event markers arising from a “visual Flanker task” presented on e-Prime software with EEG recorded using an EGI system. Please note that this script is only meant as a simplified example of an event marker labeling script, and should only serve as a starting point for your own customized scripts.

## Acknowledgement

We thank Professor Tracy Riggins for kindly providing the childhood (BioSemi) data.

