## Supplementary material for "The Maryland Analysis of Developmental EEG (MADE) Pipeline": The online supplement of this manuscript provides a step-by-step tutorial of how to use and customize the available scripts.

An EEG experiment usually includes datasets from multiple subjects and the same analysis steps are typically applied for every subject during preprocessing. Therefore, the aim of the pipeline is to make a simple protocol to batch process raw resting-state and/or event-related EEG data, particularly from pediatric populations, in order to prepare the raw EEG signal for analyses in the time and/or frequency domains. Although the pipeline has been set up as an automated preprocessing stream, it is often useful to save intermediate outputs of the analysis to inspect and visualize outputs for individual data files at particular preprocessing stages. Therefore, the pipeline will save analysis results at multiple stages in user-defined folder locations if the user wishes to do so. The following section provides a step-by-step tutorial on use of the preprocessing pipeline scripts and a description of how the lines of code can be edited to create an analysis protocol for one's own data.

**User Inputs**

As discussed above, the preprocessing of EEG data involves using the same parameters and repeating the same steps for every subject. For example, sampling rate, filter-settings, voltage thresholds for artifact rejection, and epoch length will commonly be the same for every condition and subject (unless there are special reasons for these to be different). However, some features of the data file can be subject-specific. For example, subjects will have different file names and can have different trial counts (e.g. some subjects may not complete the entire experiment, which is often the case in experiments involving pediatric populations). The pipeline takes users inputs for the common parameters to be used in preprocessing and reads subject specific information from individual data files. The user inputs section of the pipeline includes 16 questions for the user that outline parameters and decisions that the user needs to make for preprocessing the data. First, the user defines the folder location of raw data and the output folder location where the processed data will be saved. Next, the user inputs necessary parameters including channel locations, anti-aliasing time offset (if any), sampling rate, filter settings, and file format of saved data. The user also needs to make certain decisions about specific options in the pipeline: down sampling, removal of electrodes (e.g., outer layer of high density nets on infants), data epoching (then the event markers of interest and epoch length), baseline correction, epoch-level voltage rejection, epoch-level and global channel interpolation, re-referencing of data, and saving interim data files at each step.

**Analysis steps**

**Step 1: Import data.** MADE uses EEGLAB plugins to import data. For example, the pipeline uses the mffmatlabio plugin (mff_import.m function) to import EGI data files that were recorded using NetStation software. Users can modify the code to import other data formats that EEGLAB supports.

```
% import data into EEGLAB
EEG = mff_import(raw_data_location, data_file_name);
```

**Step 2: Importing channel location file**. The channel location file is imported after importing the raw data file. The channel location file is imported separately after reading the raw data to make sure the electrodes locations are labeled correctly. MADE uses EEGLAB's pop_chanedit function, which accepts almost all established channel location file formats, to import the channel location file.

```
% import channel locations file
EEG = pop_chanedit(EEG, 'load',{channels_locations 'filetype'
'autodetect'});
```

**Step 3: Adjust anti-aliasing and task related time offset (EGI Specific).** EGI's GES 300 and 400 series amplifiers delay EEG by a fixed interval inherent in the anti-aliasing filters of these amplifiers (Electrical Geodesics, Inc, 2013). This interval varies for each amplifier model and sampling rate. Additionally, in the EGI system, when aligning EEG with events that are generated from stimulus presentation or digital inputs in real time, the EEG might be offset in time relative to the event markers that are aligned with real time. Additionally, when aligning the response related events, the EEG might also be offset in time relative to the event markers. As a result, these need to be adjusted during offline processing. The amount of time that needs to be added for stimulus-related offsets can be calculated by running a visual or auditory timing test specific to the stimulus presentation (e.g. E-Prime) software and separate tests are required to calculate response-related offsets. To adjust delays caused by the EGI system and/or stimulus events, a positive value must be added to the latency of each event marker to offset this delay and to adjust for response-related events, a negative value is typically required instead. In the pipeline, the users provide input if any of time offset need to be adjusted and if no adjustment is required they can choose not to do so.

```
% Adjust time offsets
filter_timeoffset   = xxx; % EGI anti-aliasing time offset
stimulus_timeoffset = xxx; % stimulus related time offset
response_timeoffset = xxx; % response related time offset

% Adjust anti-aliasing time offset
    for aafto=1:length(EEG.event)
      EEG.event(aafto).latency = EEG.event(aafto).latency +
                                 (filter_timeoffset/1000)*EEG.srate;
    end

% Adjust stimulus related time offset
    for sto=1:length(EEG.event)
        for sm=1:length(stimulus_markers)
            if strcmp(EEG.event(sto).type, stimulus_markers{sm})
               EEG.event(sto).latency = EEG.event(sto).latency +
                                        (stimulus_timeoffset/1000)*EEG.srate;
            end
        end
    end

% Adjust response related time offset
    for rto=1:length(EEG.event)
        for rm=1:length(response_markers)
            if strcmp(EEG.event(rto).type, response_markers{rm})
               EEG.event(rto).latency = EEG.event(rto).latency -
                                        (response_timeoffset/1000)*EEG.srate;
            end
        end
    end
```

**Step 4: Changing sampling rate.** Most EEG systems record data at a sampling rate ranging from 256 Hz to 1024 Hz. However, some systems record data at a higher sampling rate (e.g. Biosemi uses from 2 kHz upwards). MADE is able to process data recorded at any sampling rate. However, it is often useful to down-sample the data to speed up processing and reduce the use of RAM and disk space when saving data.

```
% change sampling rate of data
sampling_rate = xxx; % down-sample data at xxx Hz(e.g. 250 Hz)
EEG = pop_resample(EEG, sampling_rate);
```

**Step 5: Removing outer layer of the channels.** As discussed above, removing the outermost ring of electrodes is mainly relevant for infant EEG data recorded with high-density electrodes (128 and upwards). Therefore, users would indicate "1" (yes) or "0" (no) in user input section of the pipeline as to whether they would like to remove the outer ring of electrodes. If yes, they would also provide a list of channels to be removed. The default is "0" and to keep all channels. Then, preprocessing stream will remove those channels if "1" was chosen.

```
delete_outlyr = 1; % 1=yes, 0=no,

% list of channels to be removed
RemChans = {'E17' 'E38' 'E43' 'E44' 'E48' 'E49' 'E113' 'E114' 'E119'
            'E120' 'E121' 'E125' 'E126' 'E127' 'E128' 'E56' 'E63' 'E68'
            'E73' 'E81' 'E88' 'E94' 'E99' 'E107'};

    if delete_outlyr ==1
        nbchans=cell(1,EEG.nbchan);
        for i=1:EEG.nbchan
            nbchans{i}= EEG.chanlocs(i).labels;
        end

        [chans,chansidx] = ismember(RemChans, nbchans);
        RemChans_Idx = chansidx(chansidx ~= 0);

       EEG = pop_select( EEG,'nochannel', RemChans_Idx);
    end
```

**Step 6: Filtering data.** The firfilt plugin in EEGLAB implements Windowed Sinc, Parks-McClellan, and Moving Average Finite Impulse Response (FIR) filters for low-pass, high-pass, and band-pass filtering. Furthermore, you can specify the cutoff frequency, filter order, window type (e.g. rectangular, bartlett, hann, hamming, blackman, kaiser), and the filter direction. In this pipeline, we use a non-causal (zero-phase shift) Windowed Sinc FIR filter with a hamming window for separate low-pass and high-pass filtering of data. See 'Filtering the data' section above for detail description of our specific filter choice.

```
% high-pass filter
EEG = pop_firws(EEG, 'fcutoff', high_cutoff, 'ftype', 'highpass',
      'wtype', 'hamming', 'forder', hp_fl_order, 'minphase', 0);

% low-pass filter
EEG = pop_firws(EEG, 'fcutoff', low_cutoff, 'ftype', 'lowpass', 'wtype',
      'hamming', 'forder', lp_fl_order, 'minphase', 0);
```

**Step 7: Bad channel identification and removal**. MADE employs a two-step process to identify and remove the bad channel/s from further analysis. First, it uses the 'channel_properties.m' function from the FASTER EEGLAB plugin to identify bad channels, then it employs the EEGLAB function pop_select to delete the bad channels. The details of the bad channel/s identification procedure of the channel_properties function are described in Part I of the manuscript.

```
% run faster to identify bad channel/s
list_properties = channel_properties(EEG, 1:EEG.nbchan, reference_channel);
FASTbadIdx=min_z(list_properties);
FASTbadChans=find(FASTbadIdx==1);

% reject channels that are bad as identified by Faster
EEG = pop_select( EEG,'nochannel', FASTbadChans);
```

**Step 8: Prepare data for ICA**: After filtering and removing bad channels, independent component analysis (ICA) is used to identify non-neural artifacts, such as ocular artifacts (e.g. eye blinks, saccades), muscle movements (e.g., EMG artifacts), and generic noise due to displacement or loss of scalp connection of electrode/s. MADE has adopted a special approach for ICA (see Part I), which includes the preparation of data for running ICA. The preparation involves:

1. Making a copy of dataset.
2. Applying a 1 Hz high-pass filter.
3. Inserting dummy markers and segmenting data into 1-second epochs.
4. Excluding data epochs with excessive noise by applying a wide-ranging voltage threshold (-/+1000 μV) rejection (Buzzell et et al., 2017).
5. Excluding data epochs with EMG like artifacts by applying a spectral threshold (-100-30 dB) at a frequency range (20-40 Hz), in which EMG-like artifacts are known to be expressed (Goncharova et al., 2003).
6. Excluding channels that contain excessive EMG or unusually high/low amplitudes for greater than 20% of the recording.

```
% make a copy of the EEG dataset
EEG_copy=EEG;

% apply 1Hz high-pass filter
EEG_copy = pop_firws(EEG_copy, 'fcutoff', fl_cutoff, 'ftype', 'highpass',
           'wtype', 'hamming', 'forder', fl_order, 'minphase', 0);

% insert dummy event marker 1 second apart
EEG_copy=eeg_regepochs(EEG_copy,'recurrence', 1, 'limits',[0 1],
         'rmbase', [NaN], 'eventtype', '999');

% find artifaceted epochs by outlier voltage
EEG_copy = pop_eegthresh(EEG_copy,1, 1:EEG_copy.nbchan, [-1000 1000],
           EEG_copy.xmin, EEG_copy.xmax,0,0);

% find artifaceted epochs by spectral threshold
EEG_copy = pop_rejspec( EEG_copy, 1,'elecrange', 1: EEG_copy.nbchan,
           'method', 'fft', 'threshold', [-100 30] ,'freqlimits', [20
           40], 'eegplotplotallrej', 0, 'eegplotreject', 0);

% remove artifaceted epochs identified by voltage and spectral threshold
EEG_copy = eeg_rejsuperpose(EEG_copy, 1, 1, 1, 1, 1, 1, 1, 1);
reject_artifacted_epochs=EEG_copy.reject.rejglobal;
EEG_copy = pop_rejepoch(EEG_copy, reject_artifacted_epochs, 0);
```

**Step 9: Run independent component analysis (ICA).** As discussed above, MADE performs ICA on the copied dataset that has been prepared for ICA in the previous step. MADE performs extended infomax ICA using the runica algorithm as implemented in EEGLAB. After performing ICA on copied dataset, ICA weights are transferred back from copied dataset to the original dataset.

```
% run ICA
EEG_copy = pop_runica(EEG, 'icatype', 'runica', 'extended', 1, 'stop',
                      1E-7, 'interupt','off');

% copy ICA weights that would be transferred    ICA_WINV=EEG.icawinv;
ICA_SPHERE=EEG_copy.icasphere;
ICA_WEIGHTS=EEG_copy.icaweights;
ICA_CHANSIND=EEG_copy.icachansind;

% transfer the ICA weights to the original dataset
EEG.icawinv=ICA_WINV;
EEG.icasphere=ICA_SPHERE;
EEG.icaweights=ICA_WEIGHTS;
EEG.icachansind=ICA_CHANSIND;
```

**Step 10: Identifying artifact laden independent components.** Artifact laden ICs now need to be identified. For selecting artifact laden ICs, we use the modified version of the ADJUST (adjusted-ADJUST) plugin (Leach et al., under review). In the adjusted-ADJUST, the original ADJUST algorithm has been augmented to more accurately select blinks and saccades from less straightforward pediatric ICA decompositions from geodesic nets and to de-select any ICs that include an alpha peak (likely neural activity). Just as the original ADJUST plugin, the adjusted-ADJUST identifies ICs containing eye blinks, vertical eye movements, horizontal eye movements, and generic discontinuities and produces a report for each subject. The adjust report provides a brief description of the classification of ICs based on features of each of the four kinds of artifacts.

```
% run adjust to find bad IC/s
    badIC = ADJUST(EEG_copy, [datafile_name, '_adjust_report']);

% mark the bad ICs identified by ADJUST
    for ic=1:length(bad_IC)
        EEG.reject.gcompreject(1, bad_IC(ic))=1;
    end
```

**Step 11: Removing artifact laden ICs**. After running the artifact laden ICs identification procedure, the identified ICs are excluded from the original EEG data. This procedure is done on the continuous data before defining epochs-of-interest.

```
% remove artifact laden ICs from dataset
ICs_to_remove = find(EEG.reject.gcompreject);
EEG = pop_subcomp(EEG, ICs_to_remove, 0);
```

**Step 12: Epoching data.** Epoching the data refers to segmenting the data into a fixed length around an event of interest. The pop_epoch function of EEGLAB is used to create the epochs of interest. Depending on the study design and intended analysis, epochs of any length can be created. The process of creating data epochs is same for both resting-state and event-related data.

```
% epoch data
EEG = pop_epoch(EEG, event_marker, epoch_length, 'epochinfo', 'yes');
```

**Step 13: Baseline correction**. Baseline correction denotes subtracting the mean of a time period (baseline period) from an epoch. Baseline correction is done individually for each epoch. This step is optional because baseline correction is typically applied on any analysis performed at time, frequency, or time-frequency domain. However, sometimes performing baseline correction is preferred before artifact rejection based on a voltage threshold because baseline correction can stabilize abnormal fluctuations in an epoch's time series. Users have the option of baseline correction during preprocessing and if they choose to do baseline correction at this stage, they will define the time period to be used for baseline correction.

```
% should baseline be removed during preprocess = yes/no.
remove_baseline = 'no';
baseline_window = []; % in milisecond. E.g. [-1000 -500]

% remove baseline
    if strcmp(remove_baseline, 'yes')==1
        EEG = pop_rmbase(EEG, baseline_window);
    end
```

**Step 14: Rejecting epochs containing artifacts.** The most common method used to identify and reject epochs containing artifacts is the classic voltage threshold rejection. In this method, epochs with channels that exceed a voltage threshold are deemed to contain artifacts and those epochs are rejected from the analysis. The voltage threshold depends on the features of the data and researcher's own preference. For pediatric populations, more liberal thresholds tend to be employed. For example, voltage thresholds typically used in our lab are as follows: +/-150 µV for infants, +/-125 µV for children, and +/-100 µV for adults. In the MADE pipeline, we provide a modified approach to artifact rejection that allows for epoch-level interpolation of individual channels (see Buzzell et al., 2019). In this modified approach, epochs in which ocular channels (a set of frontal channels) exceed the voltage threshold are completely removed from further analysis (not interpolated). For all other non-ocular channels that exceed the voltage threshold in a particular epoch, those channels are rejected and then interpolated within that particular epoch. However, to avoid potential bias due to excessive interpolation, epochs that contain artifacts in more than 10% of channels are rejected and completely removed from further analyses (not interpolated). Users are able to select either the traditional or the modified artifact rejection method based on their preferences. For the modified approach, users can designate a set of ocular channels for voltage threshold rejection. The modified voltage threshold method is particularly useful for pediatric populations, in which a very restricted number of trials is performed per condition. This method helps to retain more trials without compromising the quality of the data.

```
% traditional voltage threshold artifact rejection
EEG = pop_eegthresh(EEG,1, [1:EEG.nbchan], [lower_thres, upper_thres],
EEG.xmin, EEG.xmax,0,0);
EEG = eeg_rejsuperpose(EEG, 1, 1, 1, 1, 1, 1, 1, 1);
EEG = pop_rejepoch(EEG, (EEG.reject.rejthresh) ,0);
badChans=EEG.reject.rejglobal;

% modified voltage threshold artifact rejection
% select an epoch (e) after running voltage threshold
EEGe = pop_selectevent(EEG, 'epoch', e, 'deleteevents', 'off',
'deleteepochs', 'on', 'invertepochs', 'off');

% find which channels are bad for this epoch
badChanNum = find(badChans(:,e)==1);
% interpolate the bad chans for this epoch
EEGe_interp = eeg_interp(EEGe, badChanNum);
```

**Step 15: Interpolating bad channels.** After artifact rejection, the bad channels that were identified by FASTER and subsequently removed from processing are interpolated using spherical spline interpolation as implemented in EEGLAB. The EEGLAB function eeg_interp.m is used to interpolate bad channels.

```
% interpolate channels
EEG = eeg_interp(EEG, channel_location);
```

**Step 16: Rereferencing**. The last step of the preprocessing is rereferencing epoched data. The pipeline is capable of computing average rereference (i.e., the average reference over all EEG channels) or rereference to selected channels.

```
% rereferencing epoch data
EEG = pop_reref(EEG, reref);
```

The goal of this preprocessing script is to clean EEG data, specifically in pediatric populations, by removing artifacts while retaining as many trials as possible. To facilitate the inspection of the quality of the cleaned data, the pipeline also outputs a report file at the end of the preprocessing describing a variety of features from processed data. After all preprocessing steps have been completed, the data is ready to be analyzed as the researcher desires (e.g., FFT, ERP, time-frequency). The final processed data files will be either EEGLAB format (.set) or MATLAB format (.mat) depending on what the researcher designated in the users input section of the pipeline.
